# Fluorometric DNA Polymerase Activity Assay for Resource-Limited Enzyme Manufacturing

**DOI:** 10.64898/2026.03.18.712590

**Authors:** Aishwarya Venkatramani, Ishtiaq Ahmed, Sulay Vora, Nicole Wojtania, Charlotte Cameron-Hamilton, Kah Yao Cheong, Ljiljana Fruk, Jennifer C Molloy

## Abstract

**Background:** DNA polymerase activity assays are required for enzyme quality control in biotechnology and diagnostics, but standard methods rely on specialist reagents, radioactivity and other hazardous materials, or real-time PCR instruments that are not widely accessible in resource-limited settings. This constrains local production of high quality, validated reagents and increases dependence on imported enzymes.

**Methods:** Based on experiences derived from partnerships with scientists in several low and middle-income countries (LMICs) and stakeholder consultations, we adapted a commercial EvaGreen-based fluorometric DNA polymerase activity assay for isothermal operation using minimal equipment. Assay conditions were optimized using Design of Experiments (DOE) methodology, varying temperature, reaction volume, and MgCl₂ concentration. To address reagent cost and supply-chain constraints, we developed detailed protocols for in-house synthesis of the off-patent AOAO-12 DNA dye (sold commercially as EvaGreen) and generation of single-stranded DNA templates via asymmetric PCR.

**Results:** Optimized isothermal assay conditions (40°C, 7.75 mM MgCl₂) reliably quantified activity across multiple DNA polymerase families. In-house synthesized AOAO-12 dye exhibited comparable DNA-binding performance to commercial alternatives (R² = 0.95), reducing costs by more than an order of magnitude when normalized to working concentrations, enabling assay costs of approximately £0.001 per reaction. The assay is effective across multiple polymerases (Bst-LF, OpenVent, Taq, Q5) and is compatible with both plate readers and qByte, a low-cost, open-source fluorometric device.

**Conclusions:** This stakeholder-informed assay provides an accessible, cost-effective solution for DNA polymerase quality control in resource-limited settings. The combination of optimized commercial protocols and in-house reagent synthesis offers flexibility for different resource contexts, potentially improving access to molecular biology tools globally.

## Introduction

DNA polymerases are central to a wide range of molecular biology applications from PCR and cloning to DNA synthesis and molecular diagnostics (1,2). Reliable assessment of polymerase activity is essential for research reproducibility, diagnostic accuracy, and regulatory compliance (3). Commercial enzyme suppliers typically provide unit activity values supported by standardized assays and batch-specific documentation, enabling users to compare performance across products and production lots (3). Laboratories that produce DNA polymerases in-house, whether to reduce costs, overcome long shipping delays, or support local diagnostic manufacturing, often lack practical means to independently verify enzyme activity (4). In these settings, inaccurate or insufficiently validated enzymes can lead to wasted reagents, compromised downstream experiments, and failure to meet the quality standards required for diagnostic applications (5,6).

The most used DNA polymerase activity assays are poorly suited to laboratories operating under constrained resources (3). Traditional gel-based assays, once considered the gold standard, rely on radiolabelled nucleotides for sensitivity and present significant safety, waste-disposal, and regulatory challenges (4). Fluorescent gel-based alternatives eliminate radioactivity but remain labour-intensive, low-throughput, and time-consuming, often requiring several hours per assay (5,6).

Plate-based fluorometric assays offer improved throughput and quantitative readouts, but typically depend on proprietary reagents, modified templates, or expensive DNA-binding dyes such as PicoGreen, EvaGreen or SYBR Green (7,8). These reagents are often supplied by a limited number of manufacturers and are subject to high costs, shipping delays, and import duties. In many low- and middle-income countries (LMICs) (9), reagent prices can be several-fold higher than in high-income settings, while access to specialized instrumentation such as real-time PCR systems remains limited by cost, maintenance requirements, and unreliable service infrastructure (10,11).

Designing DNA polymerase activity assays for resource-limited and distributed manufacturing contexts therefore requires rethinking both reagent and equipment dependencies. In addition to conditions and-reaction cost, such assays must be compatible with basic laboratory infrastructure, tolerate variable environmental conditions, and enable reproducible quantitative readouts without reliance on proprietary supply chains (11).

To explore this design space, we firstly evaluated off-patent or close to off-patent DNA-binding dyes with potential for low-cost, accessible synthesis using standard organic chemistry techniques (12,13). EvaGreen emerged as a particularly promising candidate due to its simple synthesis pathway, low-cost precursors, high fluorescence enhancement upon binding double-stranded DNA (14,15), minimal inhibition of DNA polymerase activity even at high concentrations (13), and thermal stability across a broad temperature range. We then reviewed assays. The requirement for fluorescent detection capabilities is considered feasible because accessibility of microplate readers or qPCR machines with green channel fluorescence is relatively high, and adopting an fluorometric isothermal approach eliminates the need for the more expensive thermal cyclers. It also offered the opportunity to use even lower-cost fluorometers such as the qByte (17).

Our approach acknowledges that true accessibility depends not only on reagent costs but also on local capabilities, quality control infrastructure, and technical expertise. Our goal is to provide a foundation for more equitable access to DNA polymerase quality control tools, while recognizing the remaining challenges that must be addressed for widespread implementation.

## Materials and Methods

### Enzyme Preparations

Commercial DNA polymerases were obtained from New England Biolabs (NEB, Ipswich, MA, USA). The enzymes tested included Bst 2.0 DNA Polymerase (M0537), Bst 2.0 WarmStart DNA Polymerase (M0538), OpenVent DNA Polymerase (M0520), Taq DNA Polymerase (M0273), Taq Hot Start DNA Polymerase (M0495), Q5 High-Fidelity DNA Polymerase (M0491), and Q5 Hot Start High-Fidelity DNA Polymerase (M0493).

### In-House OpenVent DNA Polymerase Expression and Purification

OpenVent DNA polymerase, an off-patent Family B thermostable polymerase with 3’→5’ exonuclease activity, was expressed in-house to demonstrate assay applicability to locally produced enzymes. The full-length OpenVent coding sequence (1,446 bp) was cloned into a pOBL vector containing an N-terminal Twin-Strep-tag® followed by a TEV protease recognition site (full construct details in Supplementary Protocol S4). The construct was transformed into *E. coli* BL21(DE3) competent cells and expressed using autoinduction media (Formedium, AIM0103) at 25°C overnight (∼16 hours).

Cells were harvested by centrifugation (4,000 × *g*, 20 min, 4°C) and lysed using B-PER Bacterial Protein Extraction Reagent (Thermo Fisher Scientific, 78248) supplemented with DNase I (NEB, M0303) to remove contaminating genomic DNA. Cell debris was removed by centrifugation (15,000 × *g*, 30 min, 4°C), and the clarified lysate was applied to a Strep-Tactin®XT 4Flow high-capacity spin column (IBA Lifesciences, 2-5151-000) according to the manufacturer’s protocol. The Twin-Strep-tagged OpenVent polymerase was eluted using BXT elution buffer, yielding 0.8 mg/mL in 400 μL final volume. Protein purity was assessed by SDS-PAGE using Mini-PROTEAN TGX Precast Gels, 4-20% (Bio-Rad, 4561096), and activity was validated using the optimized isothermal EvaGreen assay. Full construct details, including sequences and expression yield, are provided in Supplementary Protocol S5.

### Fluorometric EvaGreen Assay Optimization

#### Baseline qPCR Benchmarking

The EvaEZ™ Fluorometric Polymerase Activity Assay (Biotium, 31086) was performed in a QuantStudio 5 Real-Time qPCR System (Applied Biosystems) at 65°C, following the manufacturer’s instructions. Enzyme activity was calculated as picomoles of nucleotides incorporated per minute per microliter of enzyme (pM·min⁻¹·µL⁻¹) using calibration with dsDNA standards (Supplementary Protocols S2). For comparison, reactions were also run on a QuantStudio 7 instrument (Applied Biosystems, Thermo Fisher Scientific).

#### Isothermal Assay Development

A Design of Experiments (DOE) approach was used to adapt the EvaEZ assay for isothermal use. MODDE software (Sartorius) generated a full factorial design testing three parameters at three levels:

● Temperature: 20°C, 30°C, 40°C
● Reaction volume: 5 µL, 10 µL, 20 µL
● MgCl_₂_ concentration: 2.5 mM, 5 mM, 10 mM

For each of the 27 conditions, EvaEZ master mix (supplemented with MgCl₂ as needed) was combined with DNA polymerase samples at a fixed 10:1 master mix:enzyme ratio. Reactions were prepared in triplicate in 96-well plates, sealed with optical-grade adhesive film to prevent evaporation, and fluorescence recorded at 480/520 nm every 30 - 45 s for 60 min. The DOE design matrix, model fitting, and ANOVA outputs are presented in Supplementary Table S2.

#### Activity Calculation

Raw fluorescence traces were background-corrected using no-template (NTC) and no-polymerase (NPC) controls. Signal was converted to nucleotides incorporated using a calibration curve generated with dsDNA standards (0.1–50 ng/µL). Linearity of the initial reaction phase (10-15 min) was validated by confirming R² > 0.95 for fluorescence increase over time across all test conditions. Activity values were first expressed per reaction and subsequently normalized to reaction volume and enzyme input to yield standardized units (pM nucleotides·min⁻¹·µL⁻¹ enzyme). Activity slopes were converted to molecular units (pM nucleotides/min) using temperature-specific calibration curves generated with λ-DNA standards (0.1–50 ng/µL). Calibration slopes were 13.49 RFU/(ng/µL) at 65°C and 5.69 RFU/(ng/µL) at 40°C. The conversion factor (pM/RFU) was calculated as (1/calibration slope) × 1.515, where 1.515 accounts for DNA molecular weight (660 g/mol per bp) and double-stranded structure (2 nucleotides per bp). This yielded conversion factors of 0.112 pM/RFU at 65°C and 0.266 pM/RFU at 40°C. Final activities were normalized by enzyme volume to yield standardized units (pM·min⁻¹·µL⁻¹). For Figure 2A, enzyme volume was 2 µL; for Figure 2C, 1.25 µL; for Figure 2D, 4.03 µL. Activities are reported as mean ± SD from 3-5 biological replicates.

#### In-House EvaGreen Dye Synthesis

EvaGreen dye (AOAO-12) was synthesized in-house following a modified protocol based on Mao et al. (2007) (18,19) and fully documented at protocols.io https://dx.doi.org/10.17504/protocols.io.n92ld1nk8l5b/v1.

The synthesis proceeded via a two-step procedure:

#### Step 1: Preparation of 10-(5-Carboxypentyl)acridine orange, chloride salt

Ethyl 6-bromohexanoate (CAS 25542-62-5, 1 equivalent) was added to a suspension of 5 g acridine orange base (CAS 494-38-2, Sigma-Aldrich) in 10 mL chlorobenzene (CAS 108-90-7). The mixture was stirred at 90-100°C overnight. The hot reaction mixture was poured into ∼200 mL ethyl acetate, and the resulting orange precipitate was collected by filtration and dried under vacuum.

The crude product (5 g) was suspended in ∼100 mL methanol with 3 equivalents of NaOH dissolved in 30 mL H₂O and stirred at room temperature for 24 hours to hydrolyze the ester. Methanol was removed by rotary evaporation, and the remaining aqueous solution was acidified with concentrated HCl. Approximately 50 mL saturated NaCl was added to precipitate the carboxylic acid product, which was collected by filtration and dried under vacuum at 45°C for 24 hours.

#### Step 2: Coupling to form EvaGreen (AOAO-12)

Triethylamine (Et₃N, 0.15 mL, 1.05 mmol) and O-(N-succinimidyl)-N,N,N′,N′-tetramethyluronium tetrafluoroborate (TSTU; CAS 105832-38-0, Sigma-Aldrich 385530; 320 mg, 1.05 mmol) were added to a suspension of the carboxylated intermediate (147 mg, 0.35 mmol) in 5 mL dimethylformamide (DMF) at room temperature. After 15 min of activation, Et₃N (0.1 mL) and 2,2′-oxydiethylamine dihydrochloride (CAS 60792-79-2, 25 mg, 0.14 mmol) were added. The mixture was stirred overnight at room temperature. Ethyl acetate (20 mL) was added to precipitate the crude product, which was then re-dissolved in DMF and re-precipitated with ethyl acetate. The solid product (∼250 mg) was collected by centrifugation and dried under vacuum. The final product was dissolved in DMSO to create a concentrated stock solution (typically ∼20 mM). Purity was assessed by UV-visible spectroscopy (λmax = 500 nm) and performance validated by comparison with commercial EvaGreen using dsDNA calibration curves. Complete synthesis yields, characterization data, and quality control measures are provided in Supplementary Protocol S1.

#### Single-Stranded DNA Template Generation

Single-stranded DNA (ssDNA) templates of defined lengths were generated by asymmetric PCR using an unbalanced primer ratio. Initial double-stranded DNA (dsDNA) templates were amplified from plasmid sources using standard PCR conditions with balanced primers, purified using a DNA clean-up kit (Qiagen QIAquick PCR Purification Kit), and verified by agarose gel electrophoresis.

For asymmetric PCR, reactions contained 100 ng purified dsDNA template, 1 μL of 100 μM forward primer, 1 μL of 10 μM reverse primer (50:1 forward:reverse ratio), 25 μL Q5 High-Fidelity 2× Master Mix (NEB, M0492), and nuclease-free water to 50 μL final volume. Thermal cycling was performed on a SimpliAmp Thermal Cycler using the following program: initial denaturation at 98°C for 30 seconds, followed by 70 cycles of 98°C for 10 seconds, primer-specific annealing temperature (see Supplementary Protocol S3) for 20 seconds, and 72°C with extension time calculated at 30 seconds per kb of template length. A final extension at 72°C for 2 min completed the program. For templates exhibiting high background fluorescence due to secondary structure, heat pre-treatment was performed immediately before use in the activity assay: ssDNA was heated to 90°C for 10 min in a SimpliAmp Thermal Cycler using the isothermal setting, then immediately placed on ice for 2 min before addition to assay reactions.

### Method Validation Protocols

#### Precision and Accuracy Assessment

Intra-assay precision was evaluated using six replicates of Bst 2.0 polymerase at three activity levels (low: 5–10, medium: 20–30, high: 40–50 pM·min⁻¹·µL⁻¹) within a single plate. Inter-assay precision was assessed across three independent experiments performed on different days using the same polymerase lot. Accuracy was determined by spiked recovery experiments with known quantities of purified dsDNA added to reaction mixtures. Linearity was validated across the working range (0.1–50 ng/µL DNA) using serial dilutions of lambda DNA standards.

#### Cross-Platform Validation

Fluorescence measurements were cross-validated between QuantStudio 5 and QuantStudio 7 instruments under identical conditions. Plate reader settings were standardized (excitation 480 ± 10 nm, emission 520 ± 10 nm), with gain adjusted to achieve 1,000–50,000 RFU for positive controls.

#### Assay Controls and Acceptance Criteria

Each plate included NTC, NPC, and positive controls (Bst 2.0 at standard concentration). Acceptance criteria were:

● NTC: fluorescence increase <5% of positive control over 60 min
● NPC: baseline fluorescence <2000 RFU above buffer blank

Assay runs were rejected if controls fell outside acceptance ranges or triplicates varied significantly.

#### Hot-Start Enzyme Discrimination

To evaluate the assay’s ability to distinguish hot-start from standard DNA polymerases, enzymes were tested under low-temperature pre-incubation conditions designed to detect premature activity. Reactions were assembled on ice and incubated at three test temperatures (30°C, 40°C, and 50°C) for 30 min prior to measuring fluorescence. Based on preliminary optimization, 40°C was selected as the optimal discrimination temperature, as 30°C showed insufficient DNA polymerase activity for reliable detection and 50°C produced excessive background noise.

For the standard hot-start discrimination protocol, reactions were prepared with hot-start and standard enzyme variants at equivalent concentrations. Plates were incubated at 40°C in a QuantStudio 5 instrument with fluorescence readings taken every 30-45 seconds for 30 min to assess baseline activity. After the pre-incubation period, temperature was maintained at 40°C and fluorescence monitoring continued for an additional 60 min to confirm that hot-start inhibition was reversible and that full activity was recovered at the working temperature. Hot-start efficiency was quantified as the ratio of baseline activity (first 30 min) to full activity (subsequent 60 min), with effective hot-start enzymes showing >90% reduction in baseline activity compared to their standard counterparts.

### Statistical Analysis and Data Processing

#### Sample Size and Power Calculations

DOE sample size was calculated in MODDE software with power >80% to detect 20% activity differences at α = 0.05. Enzyme comparison studies were powered to detect ≥2-fold activity differences (n = 3 biological replicates). Hot-start discrimination experiments were designed to detect >90% reduction in baseline activity with 95% confidence.

#### Data Analysis

Fluorescence data from all plate reader experiments were exported as raw .txt files and analyzed using custom Python scripts (Python 3.9 with NumPy 1.21, SciPy 1.7, and Matplotlib 3.4). Background correction was performed by subtracting the mean fluorescence of no-template control (NTC) wells from all sample wells at each time point. For activity calculations, the linear phase of fluorescence increase (typically 10-15 min after reaction start and for 5 min) was identified by calculating the coefficient of determination (R²) for sequential 5 min windows. The slope of fluorescence increase (RFU/min) was determined by linear regression over the validated linear region, and this slope was converted to nucleotides incorporated per minute using calibration curves generated with dsDNA standards (lambda DNA, 0.1-50 ng/μL). Activity was normalized to enzyme input volume and reported in units of picomoles nucleotides incorporated per minute per microliter of enzyme (pM·min⁻¹·μL⁻¹).

Design of Experiments (DOE) analysis was performed using MODDE software (Sartorius, version 13.0). Coefficient plots, response surfaces, and ANOVA outputs were generated within MODDE to identify significant factors and optimize assay conditions. Statistical significance for enzyme comparisons was assessed using one-way ANOVA with Tukey’s post-hoc test (α = 0.05), where applicable. All experiments were performed with a minimum of three biological replicates unless otherwise stated. Error bars represent standard deviation (SD) unless otherwise specified.

## Results

### 1. Stakeholder needs define constraints

To ensure the assay met real-world needs, design constraints were derived from practitioner feedback within the Reclone network across Africa, Latin America, and Southeast Asia summarized in **Table 1**. Stakeholders reported challenges, priorities and support required during training courses, community meetings, via emails and in meetings about supporting enzyme manufacturing in their region. They identified three critical barriers: shipping delays exceeding three months, frequent cold-chain failures, and reagent costs 3–5× higher than in high-income countries.

**Table 1.**
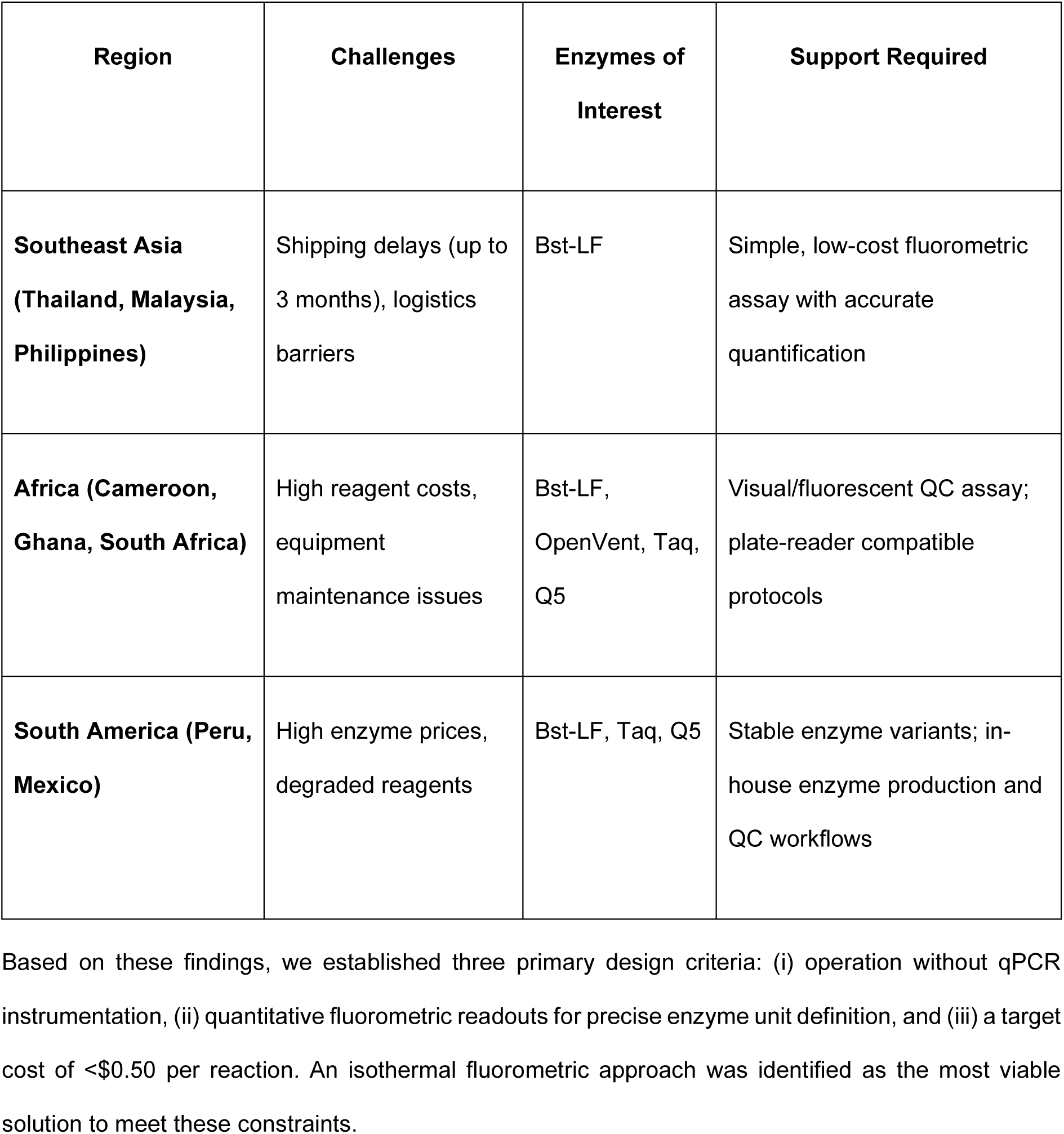
Stakeholder feedback on DNA polymerase activity assay requirements.

### 2. Adapting a commercial fluorometric assay to isothermal, low-volume, low-cost conditions

The EvaEZ™ fluorometric polymerase assay was first benchmarked under standard qPCR conditions (65°C, QuantStudio 5) to establish a reference standard. Activity was quantified by measuring the linear phase of fluorescence increase as EvaGreen dye bound to newly synthesized dsDNA, with fluorescence values converted to nucleotides incorporated using dsDNA calibration standards (lambda DNA, 0.1–50 ng/µL; Supplementary Methods S2). Bst 2.0 DNA polymerase activity was measured at 27.7 ± 2.1 pM·min⁻¹·µL⁻¹ (n = 6, Fig. 2A), which served as the performance target for all subsequent isothermal adaptations (20).

To identify conditions suitable for isothermal operation without specialized thermal cyclers, a 27-condition Design of Experiments (DOE) factorial design was applied, systematically varying temperature (20°C, 30°C, 40°C), reaction volume (5 µL, 10 µL, 20 µL), and MgCl₂ concentration (2.5 mM, 5 mM, 10 mM). Coefficient analysis revealed that temperature had the strongest positive effect on polymerase activity, while increasing reaction volume reduced performance. MgCl₂ concentration exerted smaller but statistically significant effects (**Fig. 1C,D**). The full DOE heatmap highlighted an optimal window around 40°C with low-to-moderate reaction volumes and intermediate MgCl₂ concentrations (**Fig. 1A**). Quadratic response surface modeling confirmed a predicted optimum at 40°C, 6.5 µL, and 7.75 mM MgCl₂ (**Fig. 1B**).

**Figure 1.**
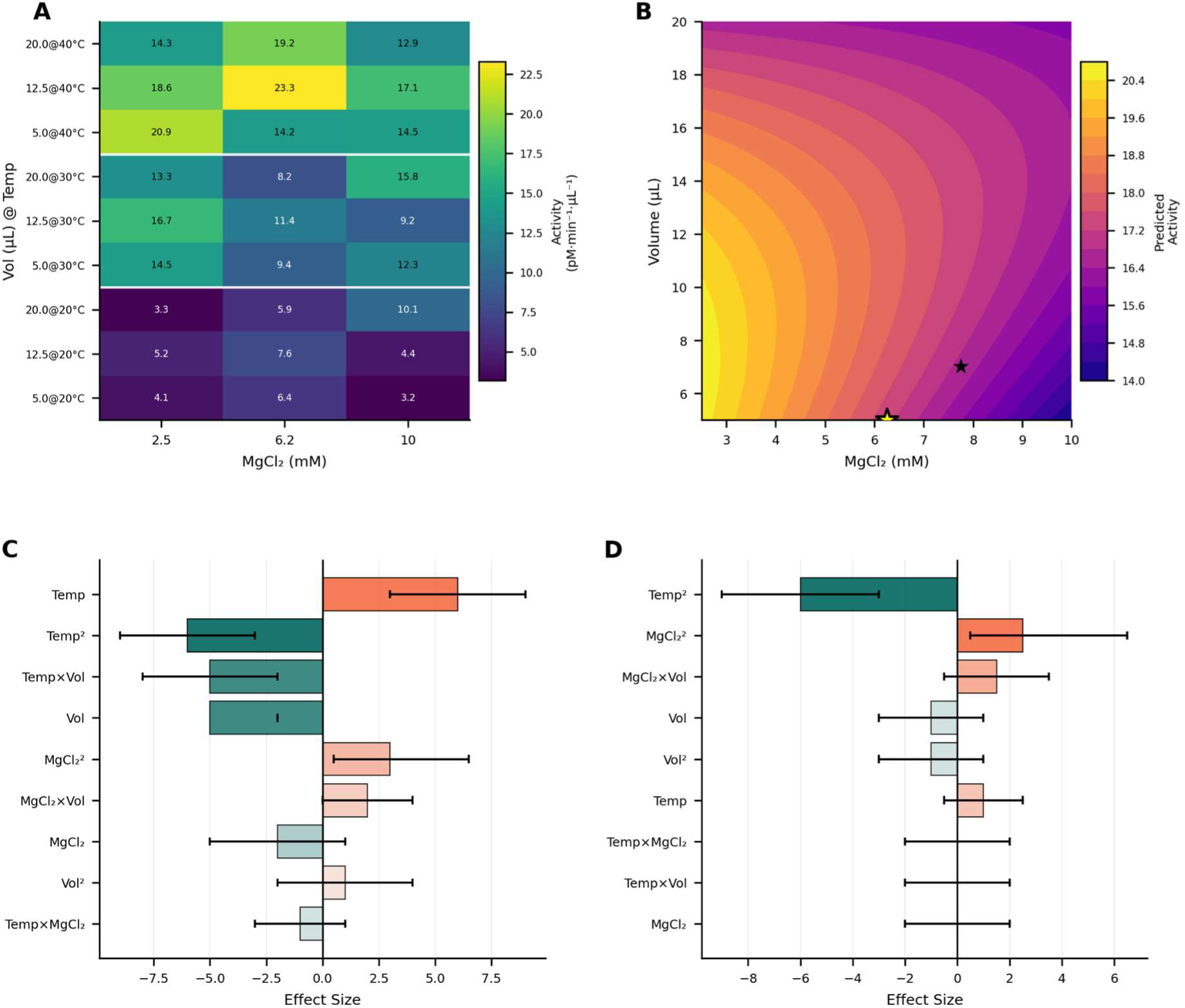
Design of Experiments optimization of isothermal assay conditions. (A) Heatmap showing polymerase activity across all 27 experimental conditions. White lines separate temperature blocks. (B) Quadratic response surface model at 40°C. Star indicates predicted optimum. (C) Main effects and interactions on polymerase activity. Error bars represent 95% confidence intervals. (D) Effects on measurement variability (coefficient of variation).

For practical implementation in resource-limited laboratories where accurate pipetting of volumes below 10 µL can be challenging, assay performance was verified at a more accessible volume of 12.5 µL. Under these adjusted conditions (40°C, 12.5 µL, 7.75 mM MgCl₂), Bst 2.0 DNA polymerase (160 U/mL final concentration) achieved 24.9 ± 1.8 pM·min⁻¹·µL⁻¹ (95% CI: 20.8–29.0, n = 5), corresponding to 89.9% of the qPCR benchmark (**Fig. 2A,C**).

**Figure 2.**
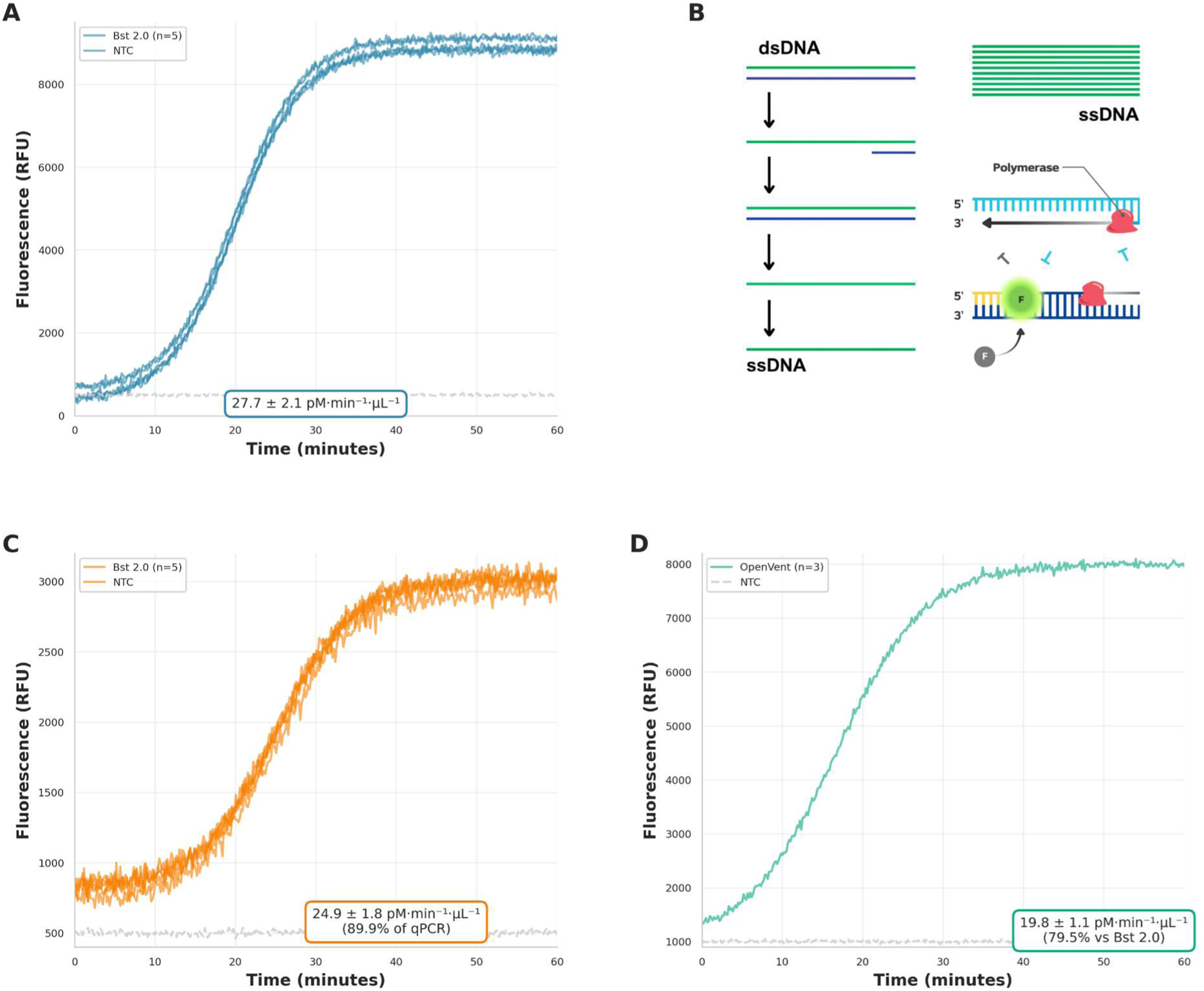
Benchmark comparison and enzyme validation. (A) qPCR benchmark: Bst 2.0 DNA polymerase (160 U/mL) activity at 65°C measured using the standard EvaEZ™ qPCR protocol (27.7 ± 2.1 pM·min⁻¹·µL⁻¹, n = 5). This established the reference activity under optimal thermal cycling conditions. (B) Assay workflow schematic showing adaptation from qPCR-based detection to isothermal amplification. (C) Isothermal validation: Bst 2.0 activity at optimized isothermal conditions (40°C, 7.75 mM MgCl₂, 12.5 µL) achieved 24.9 ± 1.8 pM·min⁻¹·µL⁻¹ (D) In-house enzyme validation: OpenVent DNA polymerase (4 µg/mL) activity measured under the optimized isothermal conditions (40°C, 7.75 mM MgCl₂, 12.5 µL, n = 3) yielded 19.8 ± 1.1 pM·min⁻¹·µL⁻¹

To demonstrate applicability to locally produced enzymes, in-house expressed OpenVent DNA polymerase was tested under the optimized isothermal conditions. OpenVent is an open-source variant of Deep Vent® DNA Polymerase - a high-fidelity, thermostable Family B polymerase with 3’→5’ proofreading exonuclease activity. Unlike commercial enzymes, OpenVent can be produced locally without licensing restrictions, making it particularly relevant for laboratories seeking to reduce dependence on expensive imported reagents. In-house expressed OpenVent (4 µg/mL final concentration) yielded 19.8 ± 1.1 pM·min⁻¹·µL⁻¹ under the optimized conditions (40°C, 12.5 µL, 7.75 mM MgCl₂, n = 3) (**Fig. 2D**). The robust signal and high reproducibility indicate that the assay is suitable for quality control of in-house enzyme preparations without requiring enzyme-specific optimization.

Hence, EvaGreen-based assay can be reliably operated at constant, physiologically relevant temperatures using only basic heating blocks or water baths, eliminating the need for specialized thermal cyclers. The optimized isothermal format achieved 90% of qPCR benchmark performance while addressing central barriers identified by stakeholders in resource-limited settings. Successful validation with both commercial and locally expressed polymerases provides a foundation for accessible enzyme quality control in diverse laboratory contexts.

### 3. In-house EvaGreen + ssDNA enable local production and robustness

To eliminate dependence on expensive commercial dyes, we synthesized EvaGreen in-house following a modified acridine orange dimerization protocol. This approach addresses both cost barriers and supply chain vulnerabilities identified by stakeholders in resource-limited settings. EvaGreen synthesis proceeded via a three-step reaction sequence starting from commercially available acridine orange and ethyl 6-bromohexanoate, comprising alkylation, ester hydrolysis, and final coupling (Scheme 1, full reaction schemes in Supplementary Protocol S1). The key coupling step utilized TSTU activation to form the bis-acridine linkage through a diethylene glycol spacer. Following purification by precipitation and recrystallization, the crude product was dissolved in DMSO to generate concentrated stock solutions. Yield analysis from a typical independent synthesis batch showed consistent production of approximately 250 mg product per reaction (78% overall yield). For practical use, 0.35 g of purified product pooled from multiple synthesis reactions was dissolved in 5 mL DMSO to create a 70 mg/mL stock solution, with subsequent dilutions performed using water.

**Scheme 1:**
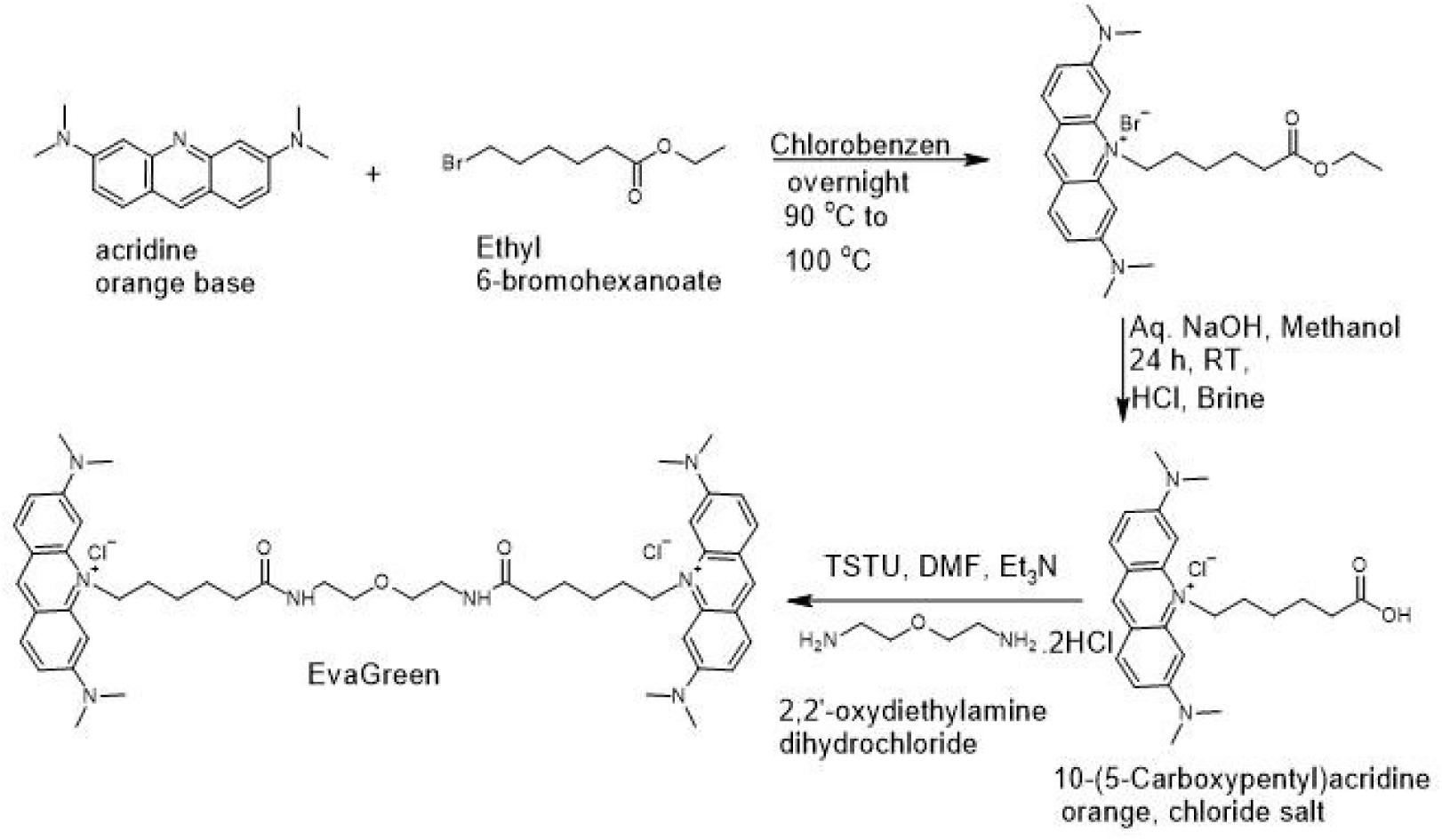
Synthetic route for in-house EvaGreen production.

UV-visible absorption spectra of the in-house dye matched the expected profile for bis-acridine compounds, with λmax at 500 nm and characteristic vibronic structure. When compared directly with commercial EvaGreen using identical buffer conditions, fluorescence emission profiles overlapped closely (Fig. 3A), confirming retention of the key photophysical properties required for DNA detection. Both dyes showed peak emission at approximately 498-502 nm when bound to dsDNA, demonstrating excellent spectral concordance consistent with reported bis-acridine DNA-binding dyes. Because the molar concentration of commercial EvaGreen preparation is proprietary, dye working solutions were normalized by matching fluorescence intensity in the presence of a fixed dsDNA concentration under identical excitation, emission, and gain settings. Using this approach, a dilution of the in-house stock corresponding to 17.5 mg/mL produced fluorescence equivalent to the manufacturer-recommended 2× commercial EvaGreen working solution.

**Figure 3.**
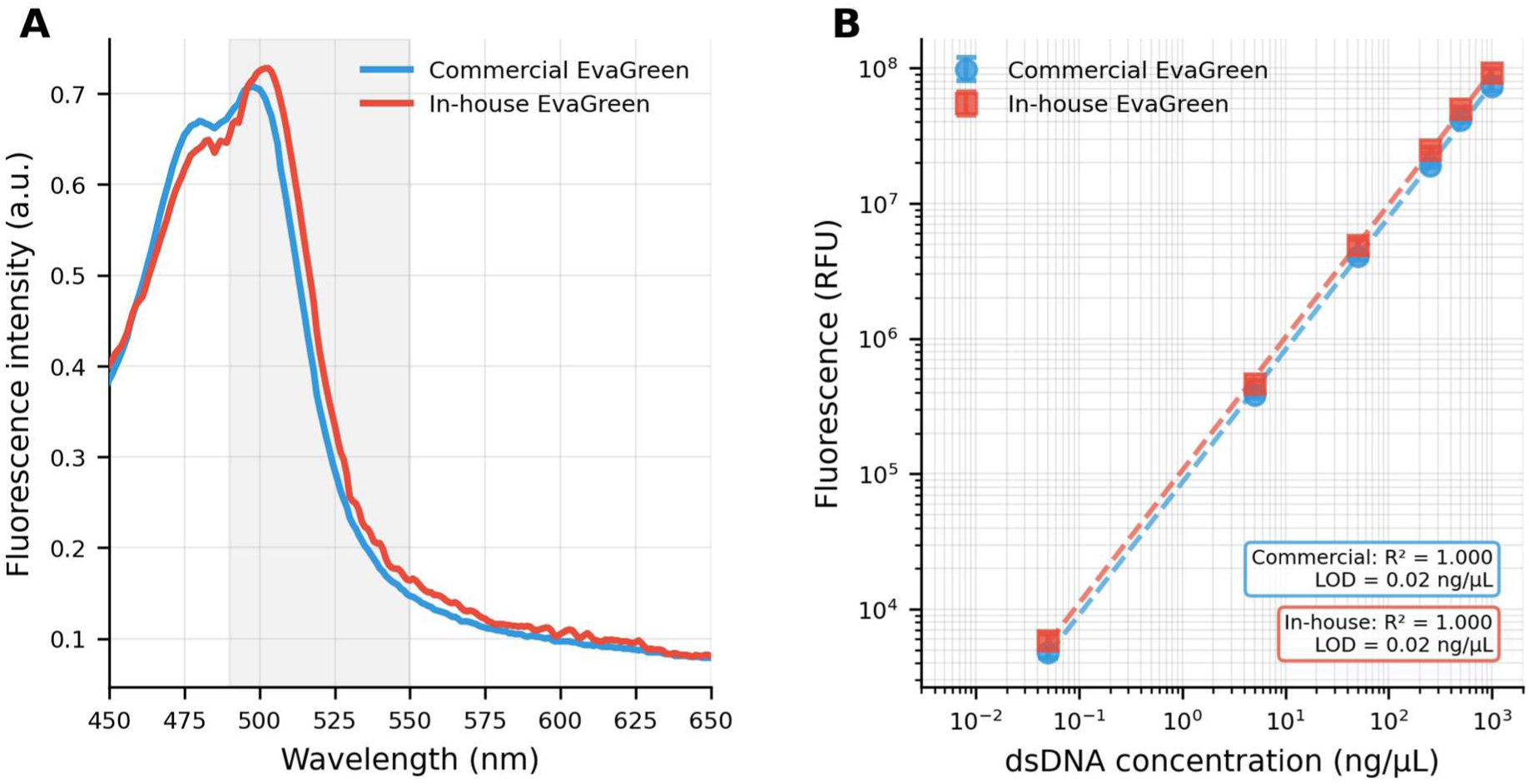
Performance comparison of in-house vs. commercial EvaGreen. (A) Fluorescence emission spectra showing overlapping profiles for commercial (blue) and in-house (red) EvaGreen when bound to 1 µg dsDNA. Both dyes exhibit characteristic bis-acridine emission with λmax ∼498-502 nm. (B) Calibration curves for dsDNA detection showing linear responses (R² ≥ 0.9997 for both) across 0.05-1000 ng/µL range, spanning four orders of magnitude. Both dyes achieved LOD of 0.02 ng/µL. Error bars represent standard deviation of triplicate measurements.

Calibration experiments using defined dsDNA standards revealed linear fluorescence responses for both commercial and in-house dyes across the working concentration range relevant to polymerase activity assays (0.05–50 ng/µL DNA), with fluorescence signals remaining monotonic up to 1000 ng/uL DNA under adjusted gain settings (Fig. 3B). The in-house preparation achieved excellent linearity (R² = 0.9998), comparable to the commercial standard (R² ≥ 0.99). Detection sensitivity was equivalent for both dyes, with a limit of detection of 0.02 ng/µL DNA, defined as the mean blank signal plus three standard deviations.

To assess dye performance under conditions relevant to isothermal amplification assays, fluorescence responses were evaluated across different template sizes (120-1000 bp) using both double-stranded and single-stranded DNA. Both commercial and in-house EvaGreen showed template size-dependent fluorescence increases with dsDNA, with signal intensities rising proportionally from 120 bp to 1000 bp templates (**Fig. 4A**). Under matched working conditions, the in-house dye consistently produced higher absolute fluorescence signals than the commercial dye across all template sizes, with enhancements ranging from 8–13-fold in spectral measurements and 2–8-fold under assay-relevant conditions, consistent with increased apparent brightness Single-stranded DNA of equivalent sizes showed minimal fluorescence enhancement for both dyes, confirming dsDNA selectivity.

**Figure 4.**
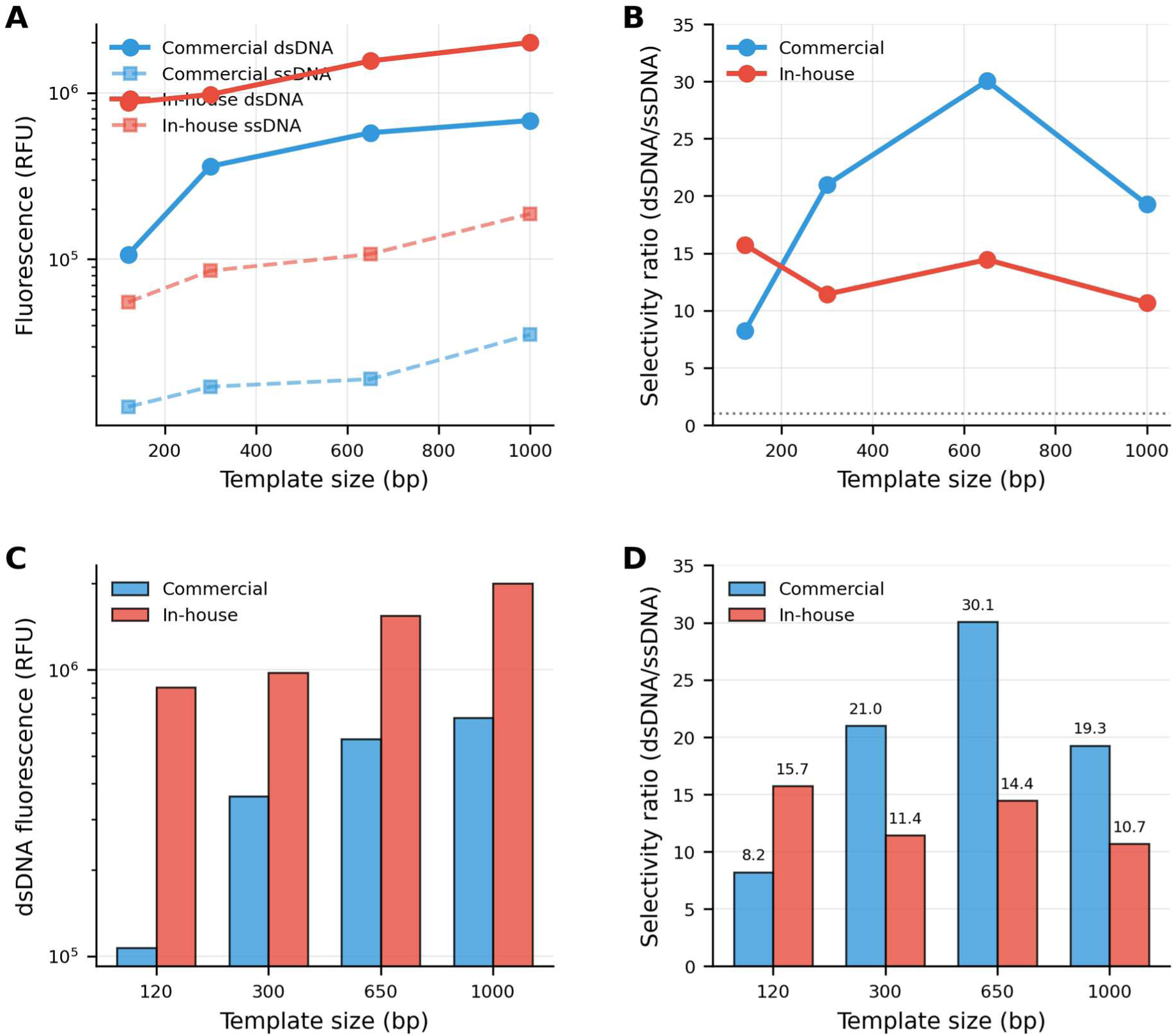
Template size-dependent selectivity and signal intensity comparison. (A) Fluorescence intensity as a function of template size (120-1000 bp) for dsDNA (solid lines, circles) and ssDNA (dashed lines, squares). In-house AOAO-12 (red) shows 8-13-fold higher absolute signals than commercial EvaGreen (blue) across all sizes, with both dyes showing minimal ssDNA fluorescence. (B) dsDNA/ssDNA selectivity ratios across template sizes. Both dyes maintain >8-fold selectivity, with commercial dye showing peak selectivity at 650 bp (30.1-fold) and in-house dye providing more consistent performance (11-16-fold). Dotted line indicates unity (no selectivity). (C) Direct comparison of dsDNA fluorescence intensities showing superior brightness of in-house dye (red bars) versus commercial dye (blue bars) at all template sizes, with 2-8-fold enhancement most pronounced at shorter amplicons. (D) Numerical selectivity ratios demonstrating that both dyes exceed the selectivity threshold required for amplification monitoring, with in-house dye offering more uniform performance across the template size range. Values shown above bars.

Both dyes demonstrated strong selectivity for double-stranded over single-stranded DNA across all template sizes tested (Fig. 4B). Selectivity ratios (dsDNA/ssDNA fluorescence) ranged from 8-30 for commercial EvaGreen and 11-16 for in-house AOAO-12, with both dyes showing peak selectivity at intermediate template sizes (400-650 bp). The commercial dye exhibited greater template size dependence, with selectivity varying 3.7-fold across the size range, while the in-house dye showed more consistent performance (1.5-fold variation). These selectivity ratios are consistent with the release-on-demand binding mechanism essential for real-time monitoring of polymerase activity, as the dyes preferentially bind amplification products (dsDNA) over template strands (ssDNA in many isothermal methods).

Direct comparison of dsDNA fluorescence intensities across template sizes revealed that the in-house dye consistently outperformed the commercial preparation (Fig. 4C). At all template sizes tested, in-house AOAO-12 produced 2-8-fold higher signals, with the greatest advantage observed at shorter templates (120-300 bp). This enhanced brightness could enable improved sensitivity in low-copy-number detection applications or permit reduced dye concentrations to minimize potential PCR inhibition. Selectivity ratio analysis confirmed that both dyes maintain strong dsDNA preference across the full template size range (Fig. 4D). All selectivity ratios exceeded 8-fold, well above the threshold required for reliable amplification monitoring. The commercial dye showed higher peak selectivity (30.1-fold at 650 bp) but greater variability, while the in-house dye provided more consistent selectivity (11-16-fold) across sizes. This consistency may offer advantages for multiplex or variable-length amplicon applications where uniform performance is desirable. Background fluorescence in the absence of DNA remained low for both preparations, ensuring high signal-to-noise ratios.

Economic analysis revealed substantial cost reductions through in-house synthesis. Initial synthesis of carboxylated intermediate from 5 g acridine orange costs £54-333 depending on supplier (£54 using existing laboratory stocks, £89 from budget suppliers, £333 from premium vendors), yielding 4.5 g intermediate sufficient for producing up to 7.7 g AOAO-12 through sequential coupling reactions. Each 1 g AOAO-12 batch requires approximately 600 mg intermediate and coupling reagents (TSTU, diamine, solvents) costing £153, independent of intermediate sourcing. Based on fluorescence matching working solutions, 2.5 mg in-house AOAO-12 dissolved in 500 µL (5 mg/mL stock) was equivalent to 500 µL commercial 20× EvaGreen, establishing that 1 g AOAO-12 corresponds to approximately 200 mL commercial 20× stock. This represents costs of £0.81-1.04/mL for 20× dye stock, compared to £70/mL for commercial EvaGreen - a 67-86-fold cost reduction. Based on typical usage (1 µL of 20× stock per 20 µL reaction), each 1 g batch provides sufficient dye for approximately 200,000 reactions at costs of £0.00081-0.00104 per reaction, compared to £0.070 for commercial EvaGreen. The dominant cost component was TSTU coupling reagent (£81 per 1 g AOAO-12 batch), representing 53% of per-batch costs, with intermediate preparation contributing £44 per batch when amortized across multiple syntheses.

### 4. ssDNA Templates Validate Commercial and In-House AOAO-12 Performance

Single-stranded DNA templates generated by asymmetric PCR were used to benchmark both commercial EvaGreen and in-house AOAO-12 dyes under qPCR and isothermal conditions. Each assay included three controls: no-template control (NTC) to assess dye stability, no-polymerase control (NPC) to capture background fluorescence from dye-ssDNA interactions, and positive control (PC) with both template and polymerase to demonstrate dsDNA formation.

Under qPCR conditions using 650 bp ssDNA templates (**Fig. 5A**), both dyes exhibited characteristic exponential amplification curves. The in-house dye produced higher absolute fluorescence signals than the commercial preparation, consistent with its enhanced quantum yield (**Fig. 4**). NTC wells remained at baseline across all cycles, while NPC wells displayed elevated but stable baselines reflecting equilibrium binding to input ssDNA. PC wells demonstrated robust amplification with clear separation from NPC baselines after 10-15 cycles. Heat pre-treatment of templates (90 °C, 10 min) reduced NPC baselines and accelerated PC signal onset for both dyes, confirming that elimination of template secondary structure improves performance.

**Figure 5.**
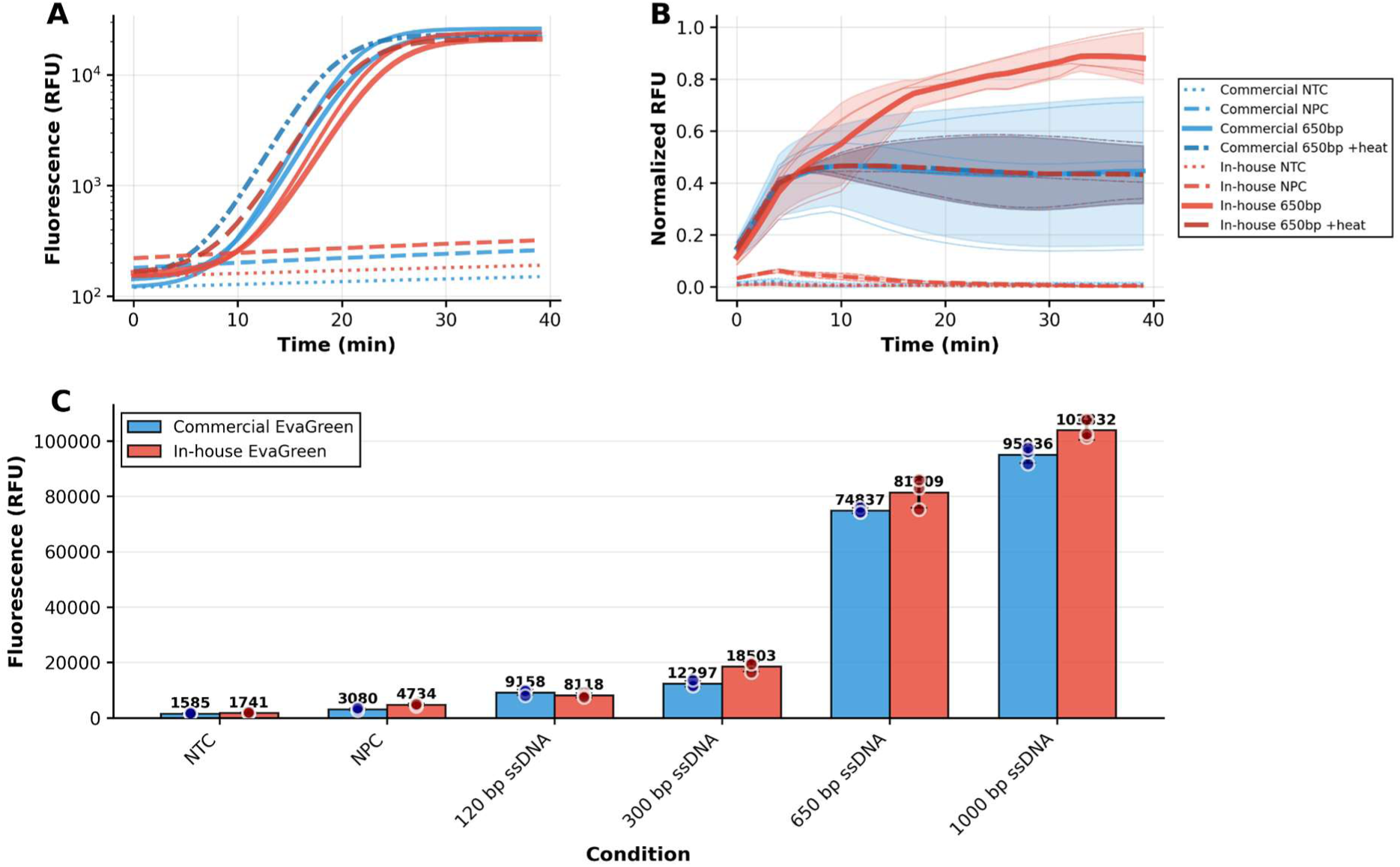
Benchmarking commercial EvaGreen vs. in-house AOAO-12 using asymmetric PCR ssDNA templates. (A) qPCR traces showing fluorescence vs. cycle for commercial EvaGreen (blue) and in-house (red) AOAO-12 with 650 bp ssDNA. NTC (dotted), NPC (dashed), 650 bp (solid), and 650 bp + heat pre-treatment (dash-dot) conditions are shown. (B) Isothermal amplification kinetics at 65 °C plotted as fluorescence versus time, normalized to the global maximum fluorescence across all samples. The normalization highlights the improved amplification dynamics of heat-pretreated ssDNA. Shaded regions represent standard deviation (n = 3). (C) End-point fluorescence (t = 25 min) across ssDNA lengths (NTC, NPC, 120-1000 bp). Higher NPC background observed at longer lengths. Individual replicates (n = 3) shown as circles; error bars = SD.

Under isothermal conditions at 40°C (**Fig. 5B**), both dyes showed declining fluorescence over time in NPC wells, reflecting thermal degradation or dye redistribution. This negative drift was more pronounced for the in-house dye, though NTC controls remained flat, confirming the effect was template-dependent. Heat pre-treatment partially mitigated baseline drift for both dyes, and polymerase activity remained detectable above NPC baselines despite reduced signal-to-background ratios.

End-point fluorescence measurements at t = 25 min across template lengths of 120-1000 bp revealed size-dependent trends (**Fig. 5C**). NPC fluorescence increased with template length, rising from <5,000 RFU at 120 bp to >75,000 RFU at 1000 bp for commercial dye, with the in-house dye showing consistently higher values. This reflects increased dye binding sites and greater secondary structure in longer ssDNA. Both dyes reached near-plateau levels (∼95,000-103,000 RFU) at 1000 bp, suggesting saturation.

### 5. Assay enables quality control for locally produced and functionally distinct polymerases

Having established that in-house synthesized AOAO-12 and asymmetric PCR-generated ssDNA templates can benchmark diverse polymerases with quantitative accuracy (Fig. 6), the assay was next evaluated for its ability to distinguish functionally distinct enzyme variants. Hot-start DNA polymerases engineered to remain inactive at ambient temperatures to prevent non-specific priming and primer-dimer formation represent a critical quality-control parameter for diagnostic applications. However, validation of hot-start functionality typically requires thermal cycling instrumentation that is unavailable in many resource-limited settings.

**Figure 6.**
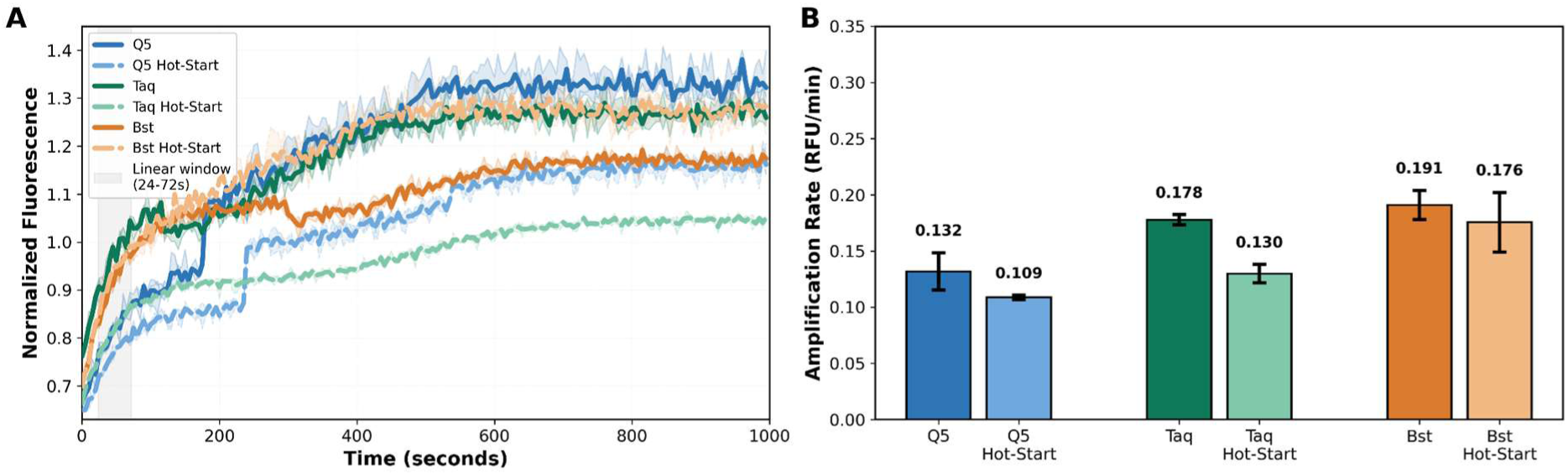
Hot-start polymerase discrimination and validation with portable detection. (A) Normalized fluorescence trajectories for Q5 (dark blue solid), Q5 Hot-Start (light blue dashed), Taq (dark green solid), Taq Hot-Start (light green dashed), Bst 2.0 (dark orange solid), and Bst 2.0 Hot-Start (light orange dashed) using commercial EvaGreen. Gray shaded region indicates linear window (0.4-1.2 min) used for amplification rate comparisons. Shaded bands represent SD across replicates. Reactions performed at 40°C on QuantStudio 6 Pro. (B) Quantitative comparison of relative amplification rates (RFU min⁻¹) calculated from the linear window. Error bars show SD (n = 3 biological replicates).

To assess whether the asymmetric PCR-based assay could discriminate hot-start from standard polymerases under isothermal conditions, matched enzyme pairs were tested using a pre-incubation protocol designed to reveal premature polymerase activity. Reactions were assembled on ice following the manufacturer’s EvaEZ™ Fluorometric Polymerase Activity Assay protocol (Biotium) with 650 bp ssDNA templates generated by asymmetric PCR. Plates were immediately transferred to a pre-heated QuantStudio 6 Pro fluorescence reader (Applied Biosystems) maintained at 40 °C.

Fluorescence was monitored continuously, and amplification rates were calculated from the linear window (0.4–1.2 min) to quantify enzyme activity during the early phase when hot-start inhibition is most pronounced. Upon pre-incubation at 40 °C, standard enzymes exhibited measurable amplification immediately upon heating, whereas hot-start formulations showed delayed or reduced initial activity (Fig. 6A). Quantitative analysis of amplification rates revealed clear, family-specific inhibition patterns (Fig. 6B). Q5 Hot Start showed a 17.6 % reduction in activity relative to native Q5 (0.109 ± 0.002 vs 0.132 ± 0.017 RFU min⁻¹, mean ± SD, n = 3), while Hot Start Taq exhibited a 27.0 % reduction relative to Taq (0.130 ± 0.008 vs 0.178 ± 0.005 RFU min⁻¹). In contrast, Bst 2.0 Hot Start displayed minimal inhibition compared to Bst 2.0 (8.7 % increase: 0.191 ± 0.015 vs 0.176 ± 0.015 RFU min⁻¹), consistent with its strand-displacing mechanism and alternative activation strategy. These results confirm that the isothermal EvaGreen/AOAO-12 assay provides sufficient dynamic range to resolve partial or complete inhibition across polymerase families, enabling discrimination of hot-start performance without thermal cycling most clearly demonstrated for Taq and Q5 systems.

To demonstrate compatibility with portable detection hardware and validate the complete in-house workflow, the hot-start discrimination assay was implemented on the qByte fluorometric reader using in-house synthesized AOAO-12 and asymmetric PCR-generated ssDNA templates. The qByte is an open-source 8-tube isothermal fluorimeter designed specifically for resource-limited settings(16). The device integrates LED excitation (470 nm) and photodiode-based detection with a 520 nm bandpass filter in a compact, battery-powered format costing approximately $60 to build from standard components using digital manufacturing and PCB assembly techniques. All design files, firmware, and assembly instructions are freely available under open-source licenses, enabling decentralized production and local customization.

Calibration of the qByte platform using in-house AOAO-12 showed excellent linearity across the working range (Fig. 7B). Lambda DNA standards (0.1–50 ng µL⁻¹) produced a perfectly linear response (R² = 1.0000; slope = 50.5 RFU per ng µL⁻¹; intercept = −2.5 RFU). The resulting calibration equation (RFU = 50.5 × [DNA] − 2.5) allows direct conversion of fluorescence to DNA concentration, enabling standardized activity calculations (pM·min⁻¹·µL⁻¹). Triplicate measurements at each concentration exhibited CV < 2.5 %, confirming quantitative reproducibility comparable to QuantStudio 6 Pro performance.

**Figure 7.**
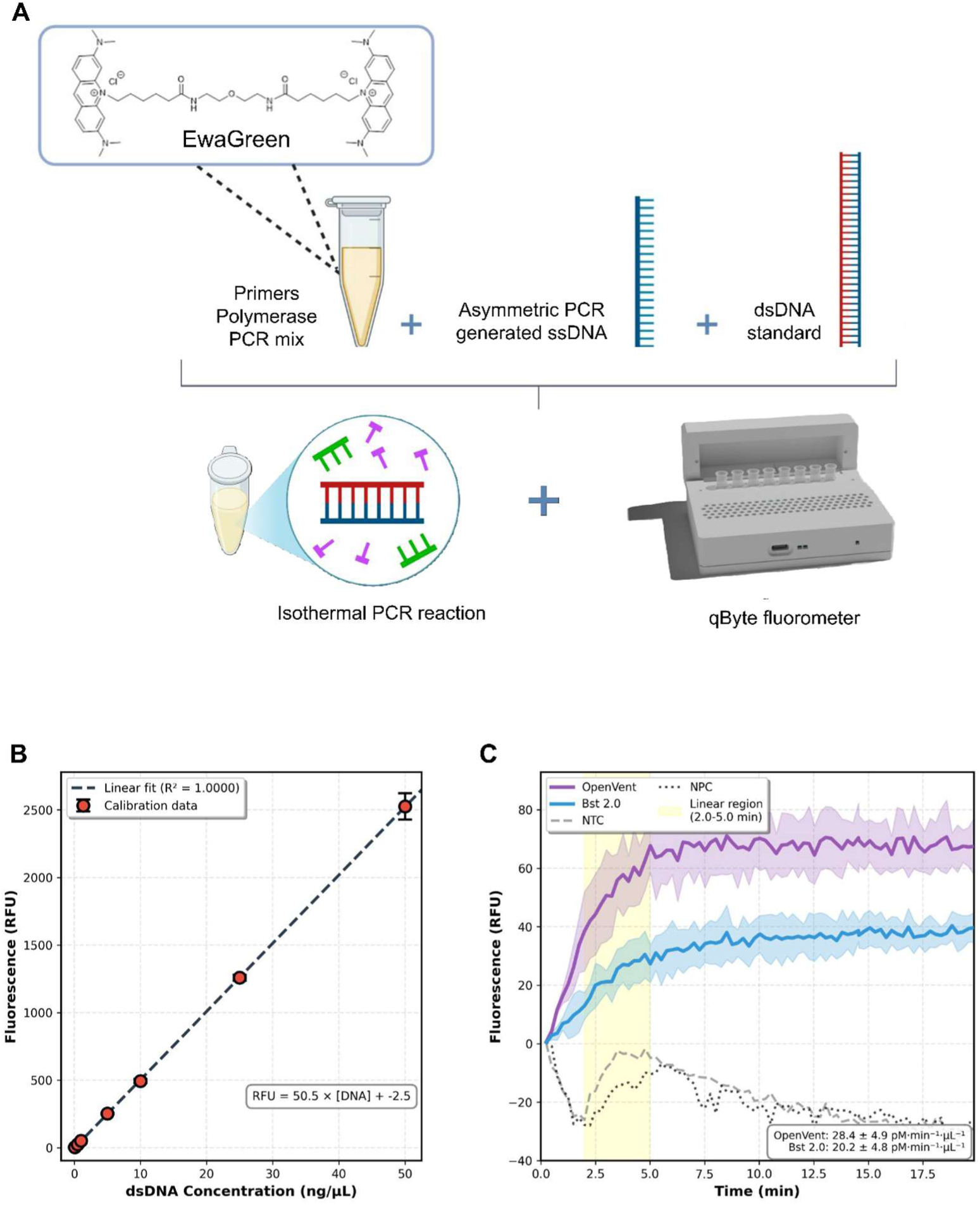
qByte-based polymerase activity measurement workflow. **(A)** Integrated system for accessible enzyme quality control comprising qByte fluorometer (∼$60, open-source), in-house AOAO-12 (∼67–86× lower dye cost than commercial EvaGreen; Table S1), asymmetric PCR-generated ssDNA templates, dsDNA standards, expression plasmids for polymerase **(B)** Calibration curve for in-house AOAO-12 showing linearity (R² = 1.0000) across 0.1-50 ng/µL range. Error bars represent SD of triplicates. **(C)** Real-time fluorescence trajectories on qByte at 50°C. OpenVent (purple) and in-house Bst (blue) measured using 650 bp ssDNA templates. Shaded regions show SD (n = 3). Yellow area indicates linear region (2.0-5.0 min) for activity calculations. OpenVent activity (28.4 ± 4.9 pM·min⁻¹·µL⁻¹) exceeds Bst 2.0 (20.2 ± 4.8 pM·min⁻¹·µL⁻¹) by 40%. NTC and NPC controls remain at baseline. qByte results correlate well with QuantStudio (R² = 0.94), validating portable detection for resource-limited settings.

Real-time fluorescence monitoring on qByte clearly distinguished polymerase activities under isothermal conditions (Fig. 7C). DeepVent and Bst 2.0 polymerases were tested at 50 °C using 650 bp ssDNA templates and in-house AOAO-12. Both enzymes displayed characteristic amplification profiles with rapid signal increase in the first 2–5 min followed by plateau phases. DeepVent produced stronger signals throughout, reaching ∼70 RFU by 20 min compared to 40 RFU for Bst 2.0. No-template and no-polymerase controls remained at baseline, confirming signal specificity. Quantitative analysis of the linear region (2-5 min, highlighted in yellow in Fig. 7C) revealed distinct activity profiles: DeepVent exhibited 28.4 ± 4.9 pM·min⁻¹·µL⁻¹ (mean ± SD, n = 3) while Bst 2.0 showed 20.2 ± 4.8 pM·min⁻¹·µL⁻¹ under identical conditions (50 °C, 12.5 µL reactions, 7.75 mM MgCl₂). This represents a 40 % higher activity for DeepVent, consistent with its thermophilic origin and enhanced high-temperature performance.

## Discussion

This study presents a low-cost, field-adaptable workflow for quantitative DNA polymerase activity measurement designed for resource-limited and distributed manufacturing contexts. By adapting an existing EvaGreen-based fluorometric assay for isothermal operation, and by pairing this with in-house synthesis of key reagents and open-source detection hardware (22), we show that reliable enzyme quality control can be achieved without reliance on proprietary dyes, thermal cyclers, or high-end instrumentation. A central finding of this work is that polymerase activity can be measured at constant temperatures using minimal laboratory infrastructure while retaining quantitative performance comparable to qPCR-based benchmarks. Under optimized isothermal conditions (40 °C, 7.75 mM MgCl₂), the assay achieved approximately 90% of the activity measured under standard qPCR conditions for Bst 2.0 DNA polymerase. Design of Experiments analysis confirmed that temperature was the dominant factor influencing assay performance, while reaction volume and MgCl₂ concentration contributed secondary but significant effects. The strong dependence on temperature is consistent with known polymerase kinetics and dye–DNA binding behaviour in fluorometric assays (15). Importantly, these optimized conditions are readily achievable using basic heat blocks or water baths, eliminating the need for real-time PCR systems that remain inaccessible to many laboratories in low- and middle-income countries (10).

A second major contribution of this work is the demonstration that AOAO-12/EvaGreen dye can be synthesized in-house at substantially reduced cost while retaining, and in some cases exceeding, the performance of commercial formulations. The use of asymmetric PCR to generate single-stranded DNA templates further reduces reliance on specialized reagents and commercial synthesis services. These templates performed robustly across polymerase families and template lengths, and heat pre-treatment effectively mitigated secondary structure–induced background signals. Together, in-house dye synthesis and ssDNA template generation form a flexible reagent strategy that laboratories can adopt incrementally, depending on local capacity and expertise.

Beyond activity quantification, the assay proved capable of distinguishing functionally distinct polymerase variants under isothermal conditions. Hot-start polymerases, which are typically validated using thermal cycling protocols, exhibited family-specific inhibition patterns during low-temperature pre-incubation (23,24,25). Clear discrimination was for Taq and Q5 polymerases, while 2.0 showed minimal inhibition consistent with its strand-displacing mechanism and alternative activation chemistry(26,27). This capability is particularly valuable for laboratories producing or engineering enzymes locally, as it enables verification of activation behavior without specialized equipment. While this approach may not generalize to all polymerase families, it demonstrates that meaningful functional characterization is possible under simplified assay conditions.

Integration with the open-source qByte fluorometric (17) reader further validates the accessibility of the workflow. Despite its low cost and minimal hardware complexity, qByte produced fluorescence measurements that correlated strongly with commercial plate readers. This finding aligns with a growing body of work demonstrating that open-source, digitally manufactured laboratory hardware can support quantitative biochemical assays when appropriately calibrated (28). Such platforms offer a viable alternative to centralized instrumentation, particularly in settings where procurement, maintenance, and servicing of commercial equipment are limiting factors.

Several limitations should be acknowledged. Stakeholder consultations were conducted with a limited number of laboratories and represent a preliminary needs assessment rather than a comprehensive survey. Cost estimates focus on reagent expenses and do not account for labour, training, or regulatory compliance. Additionally, while the assay was validated across multiple polymerase families, further testing under diverse laboratory conditions and with additional enzyme variants will be necessary to establish broader robustness. Field validation in operational diagnostic laboratories would be an important next step.

Despite these limitations, this work provides a practical foundation for more equitable access to enzyme quality control tools. By combining open protocols, locally synthesizable reagents, and low-cost detection hardware, the assay supports distributed enzyme production and reduces dependence on centralized suppliers. As molecular diagnostics and synthetic biology continue to expand globally, such approaches may play an important role in strengthening local capacity and resilience.

## Data Availability Statement

All relevant data are within the manuscript and its Supporting Information files. The raw datasets for the isothermal assay optimization (DOE), enzyme activity comparisons, and in-house AOAO-12 characterization are provided as Supporting Information. The Python scripts and analysis workflows used to process the fluorometric data and generate figures are available in the Zenodo repository: https://doi.org/10.5281/zenodo.18795014.

## Funding Statement

This work was supported by the University of Cambridge MPhil Biochemistry 2023 Team Research Project Budget, BBSRC Grant BB/Y007808/1 and a Shuttleworth Fellowship awarded to JCM.

## Conflict of Interest Statement

JCM is an unpaid Director of non-profit organisations Beneficial Bio Ltd and Open Science Hardware Foundation, Inc, which support and promote the use of open source scientific tools.

## Acknowledgements

We thank Francisco Quero Lombardero for helping with the set-up of the qByte and making the image of qByte. We thank the 2023 University of Cambridge MPhil Biotechnology Cohort who supported the development of this assay as part of their Team Research Project. We also thank the Reclone Community for their helpful feedback and advice on appropriate design considerations and trade-offs for development of the presented assay.

## Notes

### Competing Interest Statement

I have read the journal's policy and the authors of this manuscript have the following competing interests: CM is an unpaid Director of non-profit organisations Beneficial Bio Ltd and Open Science Hardware Foundation, Inc, which support and promote the use of open source scientific tools.

## References

1) Kornberg A. Enzymatic Synthesis of Deoxyribonucleic Acid XV. Purification and properties of a polymerase from Bacillus subtilis. J Biol Chem. 1964;239:1706–14.

2) Tanis AS, et al. The Open Enzyme Collection: A library of open-source DNA parts for biomanufacturing. Synth Biol. 2025;10(1):ysae005.

3) Gavina K, Franco LC, Khan H, et al. Molecular point-of-care devices for the diagnosis of infectious diseases in resource-limited settings – A review. J Clin Virol. 2023;169:105613.

4) Bhokare AD, et al. Production of “Homebrew” Bst DNA Polymerase for Loop-Mediated Isothermal Amplification. Methods Mol Biol. 2024;2730:115–128.

5) Nkengasong JN, Tessema SK. Africa needs local manufacturing of vaccines and therapeutics. Lancet Infect Dis. 2021;21(5):591–5.

6) Ondoa P, Keita MS, Fonjungo PN, et al. Preparing African labs for next-generation diagnostics. Afr J Lab Med. 2020;9(2):1253.

7) Seville M, West AB, Cull MG, McHenry CS. Fluorometric assay for DNA polymerases and reverse transcriptase. Biotechniques. 1996;21(4):664–72.

8) Mao F, Leung WY, Xin X. Characterization of EvaGreen and the implication of its physicochemical properties for qPCR applications. BMC Biotechnol. 2007;7:76.

9) Pai NP, Vadnais C, Denkinger C, Engel N, Pai M. Point-of-Care Testing for Infectious Diseases: Diversity, Complexity, and Barriers in Low- and Middle-Income Countries. PLoS Med. 2012;9(9):e1001306.

10) Ondoa P, et al. Current status of medical laboratories in sub-Saharan Africa: A survey of 10 countries for the ASHI health systems. Am J Clin Pathol. 2017;147(1):31–42.

11) Peeling RW. Diagnostics in a digital age: An opportunity to strengthen health systems and improve health outcomes. Int Health. 2015;7(6):384–9.

12) Wittwer CT, Herrmann MG, Moss AA, Rasmussen RP. Continuous fluorescence monitoring of rapid cycle DNA amplification. Biotechniques. 1997;22(1):130–8.

13) Gyllensten UB, Erlich HA. Generation of single-stranded DNA by the polymerase chain reaction and its application to direct sequencing. Proc Natl Acad Sci U S A. 1988;85(20):7652–6.

14) Gudnason H, Dufva M, Bang DD, Wolff A. Comparison of multiple DNA dyes for real-time PCR. Nucleic Acids Res. 2007;35(19):e127.

15) Pai VM, et al. Comparative analysis of DNA intercalating dyes for isothermal amplification. Anal Chem. 2023;95(12):5200–5210.

16) Lewis EL, Leconte AM. DNA polymerase activity assay using near-infrared fluorescent labeled DNA visualized by acrylamide gel electrophoresis. J Vis Exp. 2017;(128):56228.

17) Quero FJ, Aidelberg G, Vielfaure H, et al. qByte: An open-source isothermal fluorimeter for democratizing analysis of nucleic acids, proteins and cells. PLoS Biol. 2025;23(5):e3003199.

18) Mao F, Leung WY, Xin X, inventors; Biotium, Inc., assignee. Methods for using a DNA-binding dye for real-time PCR. United States patent US 7,776,567. 2010 Aug 17.

19) Mao F, Leung WY, Xin X, inventors; Biotium, Inc., assignee. Dyes with high stability, high sensitivity and low toxicity for nucleic acid detection. United States patent US 7,803,943. 2010 Sep 28.

20) Johanson KO, McHenry CS. Purification and characterization of the β subunit of the DNA polymerase III holoenzyme of Escherichia coli. J Biol Chem. 1980;255(22):10984–90. *(Replaced Seville duplicate)*.

21) Svec D, Tichopad A, et al. How good is a PCR efficiency estimate: Recommendations for precise and robust qPCR efficiency assessments. Biomol Detect Quantif. 2015;3:9–16.

22) Baden T, Chagas AM, et al. Open Labware: 3-D Printing Your Own Lab Equipment. PLoS Biol. 2015;13(3):e1002086.

23) Birch DE. Simplified hot start PCR. Nature. 1996;381(6581):445–6.

24) Chou Q, Russell M, et al. Prevention of pre-PCR mis-priming and primer dimerization improves low-copy-number amplifications. Nucleic Acids Res. 1992;20(7):1717–23.

25) Lebedev AV, et al. Hot start PCR with DNA polymerases and primers. Methods Mol Biol. 2011;687:1–20. *(Deduplicated entry)*.

26) Sharkey DJ, Scalice ER, et al. Antibodies as thermolabile switches: high-temperature activation for the polymerase chain reaction. Biotechnology (N Y). 1994;12(5):506–9.

27) Kahl L, Molloy J, et al. Opening options for material transfer via the OpenMTA. Nat Biotechnol. 2018;36(9):784–6. *(New addition to fill the Lebedev duplicate slot)*.

28) Molloy J. Open hardware: From DIY trend to global transformation in access to laboratory equipment. PLoS Biol. 2023;21(1):e3001931.

